# The NDV-3A vaccine protects mice from multidrug resistant *Candida auris* infection

**DOI:** 10.1101/465096

**Authors:** Shakti Singh, Priya Uppuluri, Abdullah Alqarihi, Hana Elhassan, Samuel French, Shawn R. Lockhart, Tom Chiller, John E. Edwards, Ashraf S. Ibrahim

**Affiliations:** Division of Infectious Disease, Los Angeles Biomedical Research Institute at Harbor-University of California, Los Angeles (UCLA) Medical Center, Torrance, California, USA; David Geffen School of Medicine, University of California, Los Angeles, CA, USA; Mycotic Diseases Branch, Center for Disease Control and Prevention, Atlanta, GA, USA

**Keywords:** *Candida auris*, NDV-3A vaccine, *Candida albicans*, Als3p homologs, biofilm, antibodies, immunization

## Abstract

*Candida auris* is an emerging, multi-drug resistant, health care-associated fungal pathogen. Its predominant prevalence in hospitals and nursing homes indicates its ability to adhere to and colonize the skin, or persist in an environment outside the host - a trait unique from other *Candida* species. Besides being associated globally with life-threatening disseminated infections, *C. auris* also poses significant clinical challenges due to its ability to adhere to polymeric surfaces and form highly drug-resistant biofilms. Here, we performed bioinformatic studies to identify the presence of adhesin proteins in *C. auris*, with sequence as well as 3-D structural homologies to the major adhesin/invasin of *C. albicans*, Als3. Anti-Als3p antibodies generated by vaccinating mice with NDV-3A (a vaccine based on the N-terminus of Als3 protein formulated with alum) recognized *C. auris in vitro*, blocked its ability to form biofilms and enhanced macrophage-mediated killing of the fungus. Furthermore, NDV-3A vaccination induced significant levels of *C. auris* cross-reactive humoral and cellular immune responses, and protected immunosuppressed mice from lethal *C. auris* disseminated infection, compared to the control alum-vaccinated mice. Finally, NDV-3A potentiated the protective efficacy of the antifungal drug micafungin, against *C. auris* candidemia. Identification of Als3-like adhesins in *C. auris* makes it a target for immunotherapeutic strategies using NDV-3A, a vaccine with known efficacy against other *Candida* species and safety as well as efficacy in clinical trials. Considering that *C. auris* can be resistant to almost all classes of antifungal drugs, such an approach has profound clinical relevance.

**Author Summary:** *Candida auris* has emerged as a major health concern to hospitalized patients and nursing home subjects. *C. auris* strains display multidrug resistance to current antifungal therapy and cause lethal infections. We have determined that *C. auris* harbors homologs of *C. albicans* Als cell surface proteins. The *C. albicans* NDV-3A vaccine, harboring the N-terminus of Als3p formulated with alum, generates cross-reactive antibodies against *C. auris* clinical isolates and protects neutropenic mice from hematogenously disseminated *C. auris* infection. Importantly, the NDV-3A vaccine displays an additive protective effect in neutropenic mice when combined with micafungin. Due to its proven safety and efficacy in humans against *C. albicans* infection, our studies support the expedited testing of the NDV-3A vaccine against *C. auris* in future clinical trials.

## Introduction

The fungus *Candida auris* was first detected in 2009 from an ear canal infection in Japan [1]. However, the earliest known strain of *C. auris* dates back to 1996 isolated in a retrospective analysis of previously misdiagnosed samples from Korea [2]. Since then, *C. auris* has been reported in more than 20 countries, with a significant number of cases detected in the Unites States [3]. Patients can remain colonized with *C. auris* for a long time and the yeast can persist on surfaces in healthcare environments, which results in spread of the organism between patients in healthcare facilities [4, 5]. Clinical isolates of *C. auris* have been recovered from a variety of specimen types, including normally sterile body fluids, wounds, mucocutaneous surfaces, and skin [4]. However, bloodstream infection remains the most commonly observed clinical manifestation of *C. auris*, with alarming in-hospital global crude mortality rates of 30 to 60% [6, 7]. Further and of high importance, some isolates of *C. auris* exhibit multidrug resistance with elevated MICs to all three major antifungal classes, including azoles, echinocandins, and polyenes, resulting in limited treatment options [8]. The ability of *C. auris* to persist and survive in an environment outside the host is unique from most other pathogenic *Candida* species. This characteristic of survival in a hostile environment perhaps means that *C. auris* has virulence determinants that help it adapt, adhere, and persist in those settings. Recent reports reveal that *C. auris* can adhere to polymeric surfaces and form highly drug resistant biofilms [5, 9, 10]. In fact, similar to other *Candida* infections, presence of a central venous catheter has been identified as a risk factor also for *C. auris* [4, 11].

We embarked on a study to determine if *C. auris* possessed evolutionarily conserved adhesin protein homologs similar to those found in another phylogenetically-related human fungal pathogen *C. albicans.* Using bioinformatic and structural homology modeling, we discovered that *C. auris* contained protein homologs of the *C. albicans* Als3p. Als3p, a member of the agglutinin-like sequence (Als) family of proteins, is also known to be conserved in other non-albicans *Candida* species [12-14], and functions as a multifunctional adhesin and invasin essential for host pathogenesis [15, 16]. The N-terminus of *C. albicans* Als3p has been developed as a vaccine that induces protective antibody and cell-mediated immune responses [17, 18], and has been shown to be safe and efficacious in clinical trials against recurrent vulvovaginal candidiasis [17, 19-24]. We found that NDV-3A vaccination induced *C. auris* cross-reactive antibody and T cell responses. Moreover, the sera from NDV-3A vaccinated mice blocked virulence characteristics of *C. auris* and enhanced opsonophagocytic killing of this fungus. Furthermore, vaccination with NDV-3A antigen protected mice from lethal *C. auris* infection and synergized with antifungal drugs. These findings identify NDV-3A as a promising vaccine for adjunctive treatment of life-threatening bloodstream infections caused by the multidrug resistant *C. auris*.

## Results

### *C. auris* possesses homologs of *C. albicans* Als3

Bioinformatic and 3-D structural modeling strategies identified three conserved proteins in *C. auris* with homologies to *C. albicans* Als3p. In particular, these *C. auris* Als3p homologs, PIS50650.1, PIS50263.1 and XP_018167572.2, displayed ~30% identity and ~50% similarity (with several regions of homology reaching up to 80-100%) to *C. albicans* Als3p. Furthermore, sequence alignments revealed that all three proteins possessed domains characteristic of Als3p, such as an N-terminal secretory signal sequence, central amyloid forming and repeated Ser/Thr-rich sequences, as well as the presence of a C-terminus GPI anchor. (Table 1, Figure S1-3). Once identified, the above sequences were assembled to produce 3-D structural models for analysis in relation to Als3p (Swiss-model figures shown in Figure 1). Based on this modeling strategy, *C. albicans* Als3p shares striking similarity to the *C. auris* proteins, particularly in the N-terminus motif (Figure 1 and S1-3).

**Figure 1.**
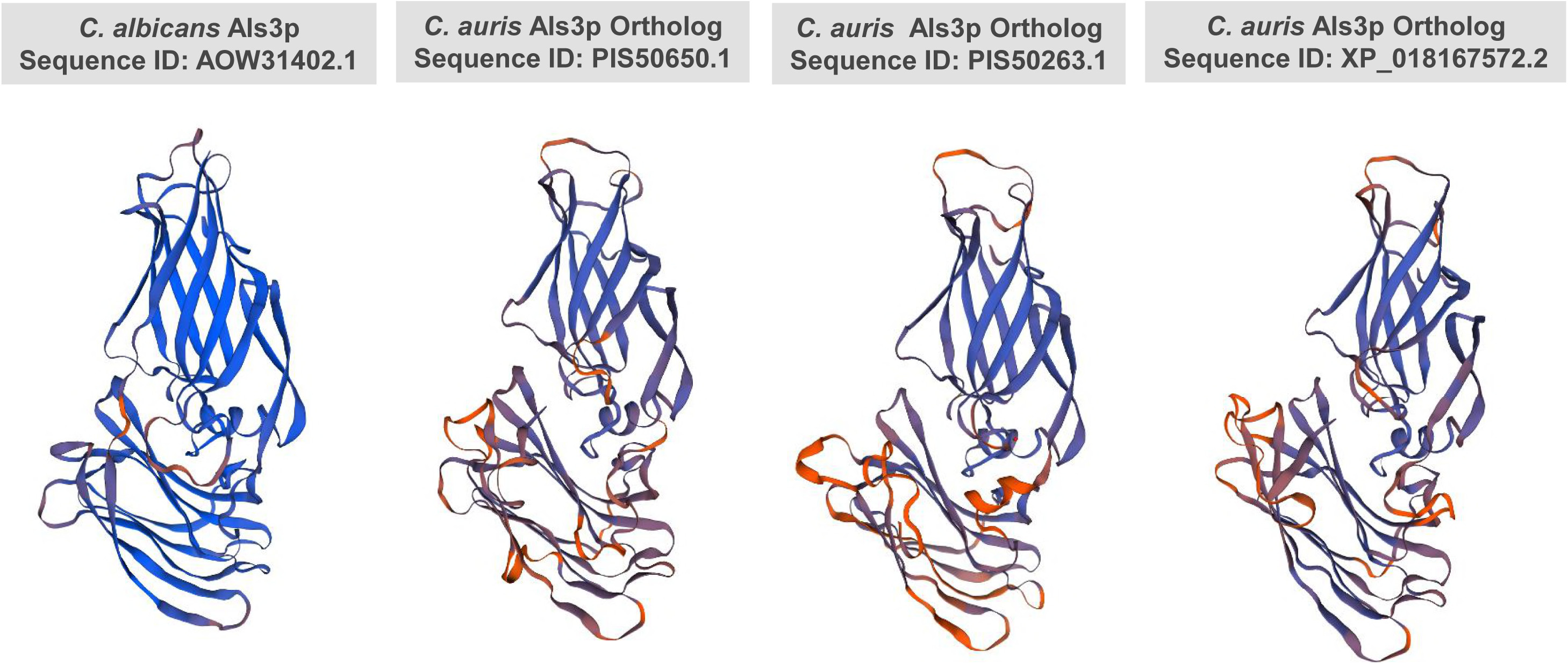
Similarity between 3-D structural models of *C. albicans* Als3p and *C. auris* Als3p homologs. Template were searched with BLAST and HHBlits was performed against the SWISS-MODEL template library (SMTL). Models were built based on the target-template alignment using ProMod3. Coordinates which was conserved between the target and the template were copied from the template to the model. Insertions and deletions were re-modelled using a fragment library. Side chains were then rebuilt. The geometry of the resulting model is regularized by using a force field. The models with high accuracy values were selected and are depicted for comparison.

**Table 1.** List of the *Candida albicans* [SC5314] Als3p orthologs in *Candida auris* with percent homology and structural features.

| S. No. | NCBI Protein Database Sequence ID | Name | Length | Match | Range | Identities | Positives | GPI-Anchor Location | Ser/Thre Rich Repeats | Amyloid Sequences |
| --- | --- | --- | --- | --- | --- | --- | --- | --- | --- | --- |
|  | AOW31402.1 | Als3p [*Candida albicans* SC5314] | 1155 | - | - | - | - | C-23 | Yes | Yes |
| 1 | PIS50650.1 | Hypothetical protein CJI97_004175 [*Candida auris*] | 1804 | 1 | 1 to 371 | 124/381(33%) | 187/381(49%) | C-23 | Yes | Yes |
|  |  |  |  | 2 | 951 to 1471 | 196/554(35%) | 287/554(51%) | C-23 | Yes | Yes |
| 2 | PIS50263.1 | Hypothetical protein CJI97_004954 [*Candida auris*] | 835 | 1 | 2 to 348 | 111/356(31%) | 179/356(50%) | C-17 | Yes | Yes |
| 3 | XP_018167572.2 | Hypothetical protein QG37_06265 [*Candida auris*] | 861 | 1 | 8 to 431 | 115/435(26%) | 204/435(46%) | C-21 | Yes | Yes |

**Note:** Protein amino acid homology was analyzed by NCBI BLAST, GPI-anchor location was predicted by GPI-SOM tool (http://gpi.unibe.ch/), Serine or Threonine rich repeats were detected by using RADAR tool (https://www.ebi.ac.uk/Tools/pfa/radar/), and amyloid sequences were detected by PASTA 2.0 tool (http://protein.bio.unipd.it/pasta2/).

### Anti-Als3p antisera from NDV-3A-vaccinated mice recognize *C. auris*

Because bioinformatic analysis revealed considerable sequence as well as 3-D structural homology between *C. albicans* Als3p and homologous proteins of *C. auris*, and predicted cell surface localization of these homologs, we hypothesized that anti-Als3p antibodies should recognize *C. auris in vitro*. The anti-Als3p antibodies were generated by vaccination of mice with NDV-3A, a vaccine based on the N-terminus of *C. albicans* Als3p, which is known to induce high titers of serum anti-Als3p antibodies [18, 20, 24]. Sera from these alum-vaccinated mice were examined in two different binding assays *in vitro*. First, different *C. auris* clinical isolates (CAU-01, 03, 05, 07 and 09) and *C. albicans* cells were grown for 90 min under germ-tube inducing conditions since Als3p is expressed on *C. albicans* hyphae [25], and then treated with anti-NDV-3A sera, followed by immunostaining with fluorescent labeled secondary IgG. Immunostaining of germinating *C. albicans* cells (positive control), confirmed the specificity of Als3p to the *C. albicans* filaments. *C. auris* yeast cells do not differentiate into hyphae, yet anti-Als3p antibodies bound to the cell surface of the fungus, as depicted by a diffused or punctate green fluorescence of the yeast cells. Fungal cells treated with sera from alum-vaccinated mice failed to fluoresce (Figure 2A). The binding of anti-Als3p antibodies to five strains of *C. auris* obtained from different clades showed the universal presence of Als3p homologs in this multidrug resistant yeast. Fungal cells treated with sera from alum-vaccinated mice failed to fluoresce (Figure 2A).

**Figure 2.**
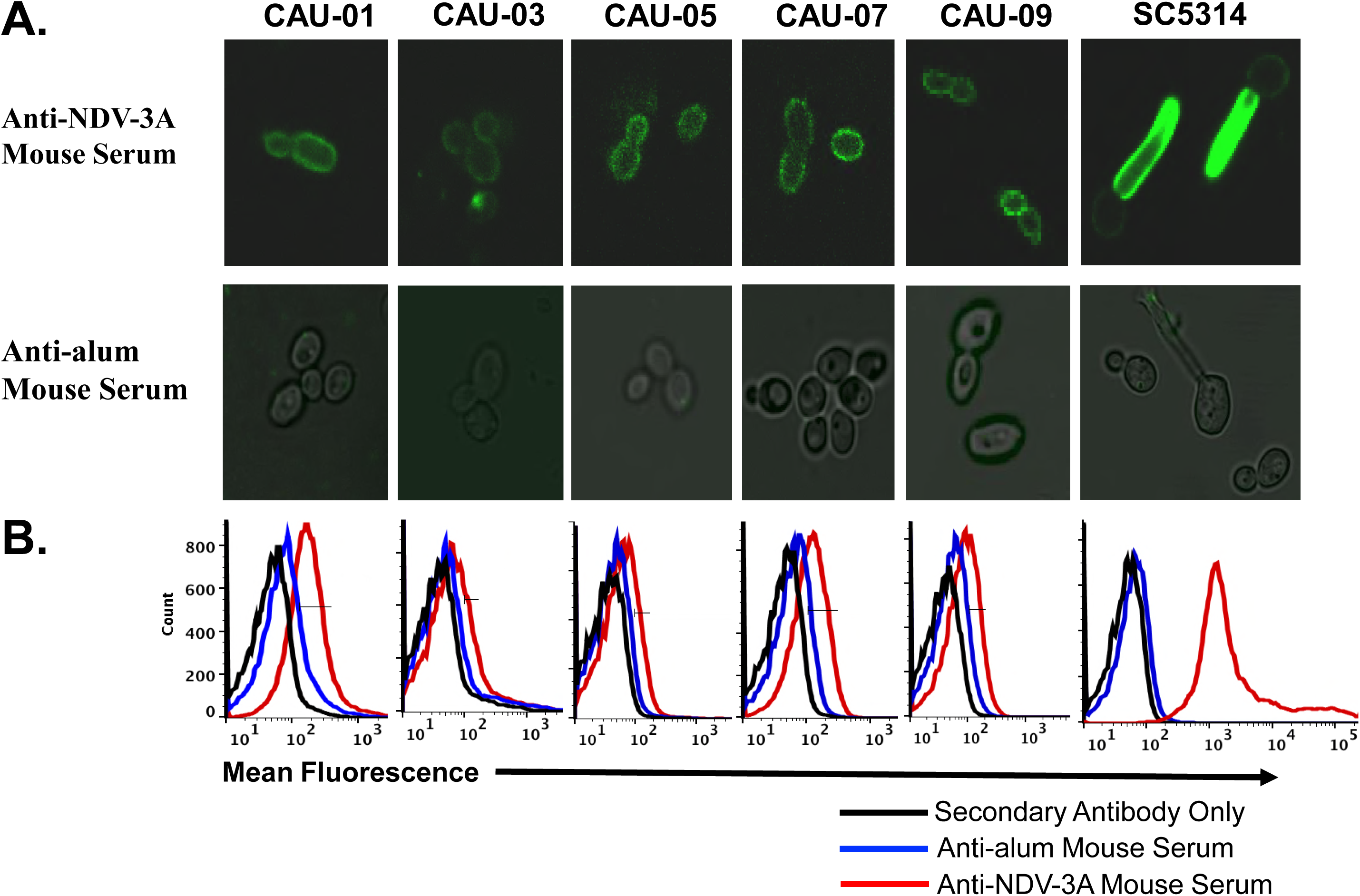
Binding of anti-Als3p IgG antibodies to *C. auris* surface. Binding of mouse IgG from NDV-3A- or alum-immunized mice with *C. albicans* (SC5314) and *C. auris* isolates (CAU-01, 03, 05, 07 and 09). The binding was detected with Alexa Fluor 488 labelled anti-mouse IgG secondary antibodies. **A)** In confocal microscopy, green fluorescence indicates binding of anti-Als3p IgG on cell surface. **B)** The stained cells acquired in flow cytometry and overlay histogram was made. The red line in the histogram shows the binding levels of anti-als3 antibodies on the cell surface compared to control mouse serum (alum).

Next, the extent of anti-Als3p antibody binding to the fungal cells was quantified by flow cytometry. The anti-Als3p antibodies present in the NDV-3A-vaccinated sera bound significantly to *C. auris* isolates (indicated by a shift in the peak representing mean fluorescence intensity), while the negative control did not. These results demonstrate the specificity of anti-Als3p antibodies to *C. auris* and further validate the modeling strategy which previously revealed the homology of *C. albicans* Als3p to Als-like proteins on *C. auris* (Figure 2B).

### NDV-3A vaccination induced robust *C. auris* cross-reactive antibodies and T-cell responses

We have previously shown that NDV-3A vaccine mediated protection against *C. albicans* infection required both Als3p-specific antibodies and CD4+ Th1/Th17 immune responses, and anti-Als3p antibody titer threshold predicts protective efficacy [21]. Thus, we reasoned that robust antibody and T cells responses would be critical for protection against *C. auris in vivo*. Therefore, we examined the magnitude of anti-Als3p antibodies binding to *C. auris* antigens by using cell-based ELISA. First, we confirmed that NDV-3A-vaccinated sera harbored high levels of anti-Als3p antibodies, by using ELISA coated with recombinant Als3p N-terminus. As expected, NDV-3A-vaccinated mice sera contained high titers of anti-Als3p antibodies (mean end-point titer of 12,800), while alum vaccinated sera had none. Next, we developed a cell-based ELISA of *C. albicans* or *C. auris* to compare the magnitude of anti-Als3p antibodies to antigens of yeast cells. We allowed the binding of serially diluted mouse sera to *C. albicans* germ tube cells or *C. auris* yeast cells and detected the endpoint titer with HRP-conjugated anti-mouse IgG detection antibodies. We found that NDV-3A vaccinated mice had high titers of cross-reacting antibodies against *C. albicans* cells, with titers similar to the recombinant Als3p N-terminus-based ELISA (12,800). Consistent with the confocal microscopy and flow cytometry results, and compared to sera obtained from alum-vaccinated mice, NDV-3A-vaccinated mice sera were found to have cross-reactive anti-Als3p antibodies to *C. auris* cells albeit with 4-fold less than *C. albicans* (mean end-point titer of 3,200) (Figure 3A).

**Figure 3.**
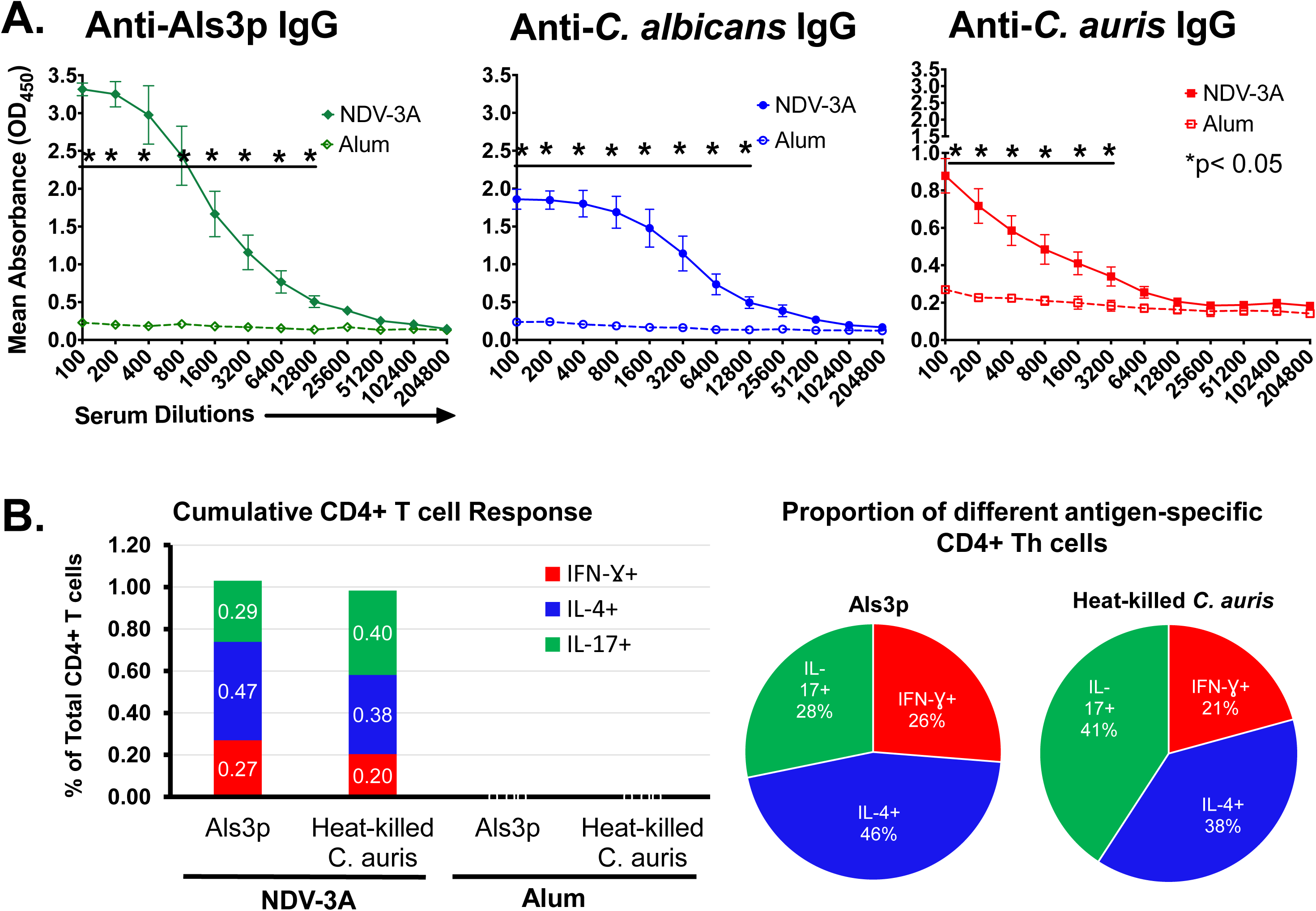
Evaluation of *C. auris* cross-reactive antibodies and CD4+ T cells in NDV-3A-vaccinated mice. **A)** Sera samples from NDV-3A-vaccinated mice were used to detect anti-Als3p antibodies in antigen ELISA and anti-*C*. *albicans* and anti-*C. auris* IgG antibodies by using cell-based ELISA. Serially diluted serum samples were incubated in Als3p-coated 96-well plate or *C. albicans*, or *C. auris* whole cell lysate as antigens. The OD450 values for each mouse were plotted against the serum dilution and mean endpoint titers were determined against each antigen by twoway ANOVA between NDV-3A and alum control for each dilution. **B)** The splenocytes from individual NDV-3A- or alum-vaccinated mice (n=4) were stimulated for 5 days with no antigen, Als3p or heat-killed *C. auris* as recall antigens. After 3 hours stimulation with PMA/ionomycin in presence of protein transport inhibitor, the cells were stained for CD3, CD4, IFN-γ, IL-4 and IL-17. The *%* of Th1 (IFN-γ+), Th2 (IL-4+) and Th17 (IL-17+) CD4+ T cells were reported after subtracting the no antigen stimulation values. Data is reported as mean of total CD4+ T cell responses (Left panel). Pie chart on the right shows proportions of antigen specific T helper cell profile.

We also examined whether NDV-3A vaccination induced memory CD4+ T cells that are also cross-reactive to *C. auris*. Splenocytes from NDV-3A- or alum-vaccinated mice were stimulated with either recombinant Als3p-N-terminus or heat-killed *C. auris* for 5 days, followed by treatment with PMA/ionomycin in the presence of monesin/brefeldin A protein transport inhibitor. These cells were stained with fluorescent-labelled antibodies against surface markers and intracellular cytokines. Flow cytometry analysis of these stained cells revealed a significant presence of high frequency of Als3p-specific and *C. auris* cross-reactive CD4+ T cells, which were comprised of Th1 (IFN-γ +), Th2 (IL-4+) and Th17 (IL-17+) cells (Figures 3B, S5-S8). The *C. auris* cross-reactive T cells were not detected in alum control mice. Further, the total *C. auris* or Als3p-specific CD4+ T cell responses were similar in magnitude and were slightly biased towards Th2 type. Finally, the *C. auris* -specific Th17 cell response was higher compared to Als3p-specific responses (Figures 3B, S5-8). Overall, these results revealed that, NDV-3A induced robust *C. auris* cross-reactive antibodies as well as CD4+ T cell immune responses.

### Anti-NDV-3A sera reduced *C. auris* biofilm formation and enhanced opsonophagocytic killing of *C. auris* by macrophages

Attachment of microorganisms to an abiotic surface is the first step in formation of a drug-resistant biofilm. Biofilm formation has been most studied in *C. albicans* and Als3p is essential for establishment of the early stages of biofilm growth [26]. Recently, we have reported that blocking Als3p by anti-Als3p antibodies abrogate *C. albicans* biofilm formation [20]. *C. auris* also has the capacity to form drug-resistant biofilms [9]. Thereby, we evaluated if anti-Als3p antibodies from NDV-3A-vaccinated sera could similarly inhibit biofilm formation in *C. auris*. Indeed, we found that sera from NDV-3A-vaccinated mice significantly inhibited biofilm formation compared to control anti-alum sera (*p* =0.001) (Figure 4A). Of note, sera were used at 1:10 (sera to media) ratio, and heat-treated prior to their contact with fungal cells, to rule out the role of complement in the inhibitory activity.

**Figure 4.**
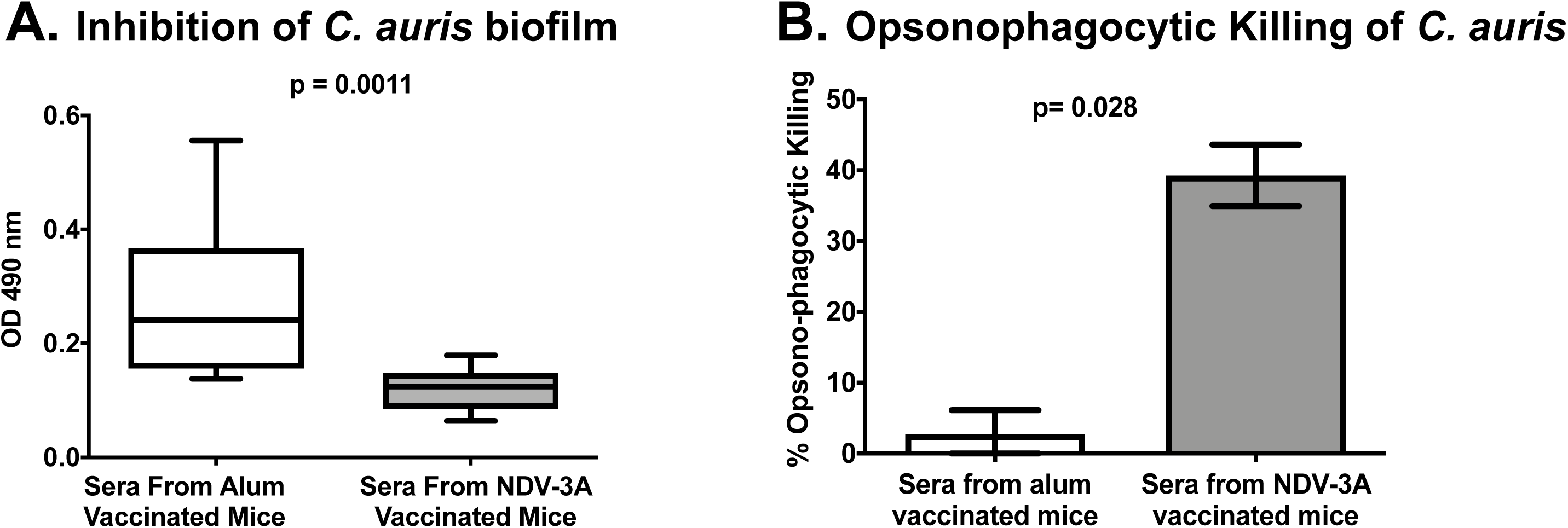
Anti-Als3p sera inhibit *C. auris* biofilm formation and augment OPK of the yeast by murine macrophages. **A)** *C. auris* biofilm formation were evaluated in 96-well plate in presence of serum from NDV-3A- or alum-vaccinated mice. The serum from NDV-3A-vaccinated mice significantly reduced *C. auris* biofilm compared to alum control mice (n=10). **B)** OPK of *C. auris* by macrophages was evaluated in presence of serum from NDV-3A- or alum-vaccinated mice. The serum from NDV-3A-vaccinated mice significantly enhanced yeast OPK compared to alum group (n=4). Results are presented relative to macrophage killing of *C. auris* without any added serum. Statistical significance was determined by Mann-Whitney Test. P values <0.05 was considered significant.

We previously demonstrated that anti-Als3p antibodies act as an opsonin to enhance phagocyte-mediated killing of *C. albicans* [20]. Thus, we posited that Als3p antibodies would enhance sensitivity of *C. auris* to macrophage killing. To test this hypothesis, heat-inactivated mice sera (at 10% concentration) from NDV-3A- or alum-vaccinated mice were incubated with *C. auris* to allow binding of anti-Als3p antibodies to the fungal cell surface, and then added to mice macrophage cell-line (1:1). As a control, *C. auris* yeast cells were subjected to macrophages without any sera. Both NDV-3A- and alum-vaccinated mice sera did not have any negative impact on growth of *C. auris* (Figure S4). However, sera from NDV-3A-vaccinated mice significantly enhanced macrophage-mediated killing of *C. auris* compared to sera from alum vaccinated mice (40% vs 2% killing respectively, normalized to *C. auris* killing by macrophage-only condition, p=0.028) (Figure 4B). Thus, anti-Als3p antibodies generated by the NDV-3A vaccine enhanced the opsonophagocytic killing of *C. auris*.

### NDV-3A vaccination protected mice from lethal *C. auris* disseminated infection

Blood stream infections are a predominant manifestation of *C. auris*, accompanied by significant rates of mortality in susceptible patients. Since, anti-Als3p antibodies generated by NDV-3A vaccination bound to *C. auris*, blocked virulence and rendered it susceptible to macrophages, we reckoned that NDV-3A vaccination would have the potential to protect against *C. auris* disseminated candidiasis. We developed a neutropenic mouse infection model by screening different clinical isolates of *C. auris* (CAU-01, 03, 05, 07, 09) for their lethality in mice. An intravenous 5x10^7^ CFU/mouse dose of *C. auris* CAU-09 showed 100% mortality in neutropenic mice and was used for vaccine efficacy testing (immunocompetent mice are resistant to *C. auris* bloodstream infection). To test the vaccine-mediated protective efficacy, mice were either vaccinated with NDV-3A or alum, three times (primary + two booster doses), and then infected with *C. auris* via the tail vein (Figure 5A). Mice vaccinated with alum displayed 100% mortality by day 4, while NDV-3A-vaccinated mice showed 40% overall survival. Surviving mice appeared healthy at 21 days when the experiment was terminated (Figure 5B). Furthermore, while both sets of vaccinated mice displayed weight loss post infection compared to uninfected mice, NDV-3A-vaccinated mice had lesser extent of weight loss versus alum-vaccinated mice (Figure 5C panel). In replicate studies to the survival experiments, mice were sacrificed 4 days post infection, and their target organs (kidneys and brain) were collected for tissue fungal burden. The NDV-3A-vaccinated mice had a 10-fold lower (~1.0 log) fungal burden in both kidneys and brains compared to alum control mice (*p*=0.0006) (Figure 5D).

**Figure 5.**
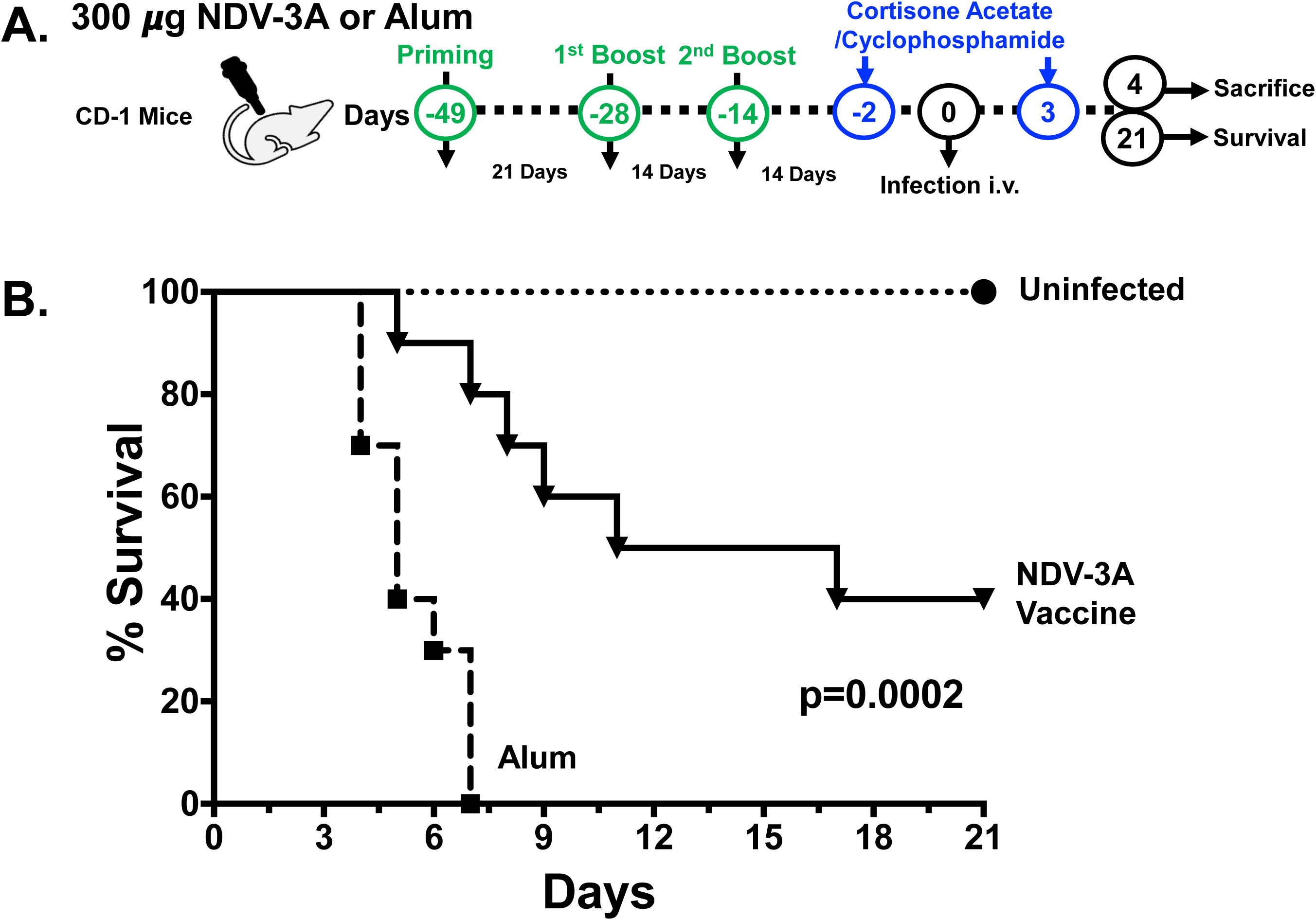
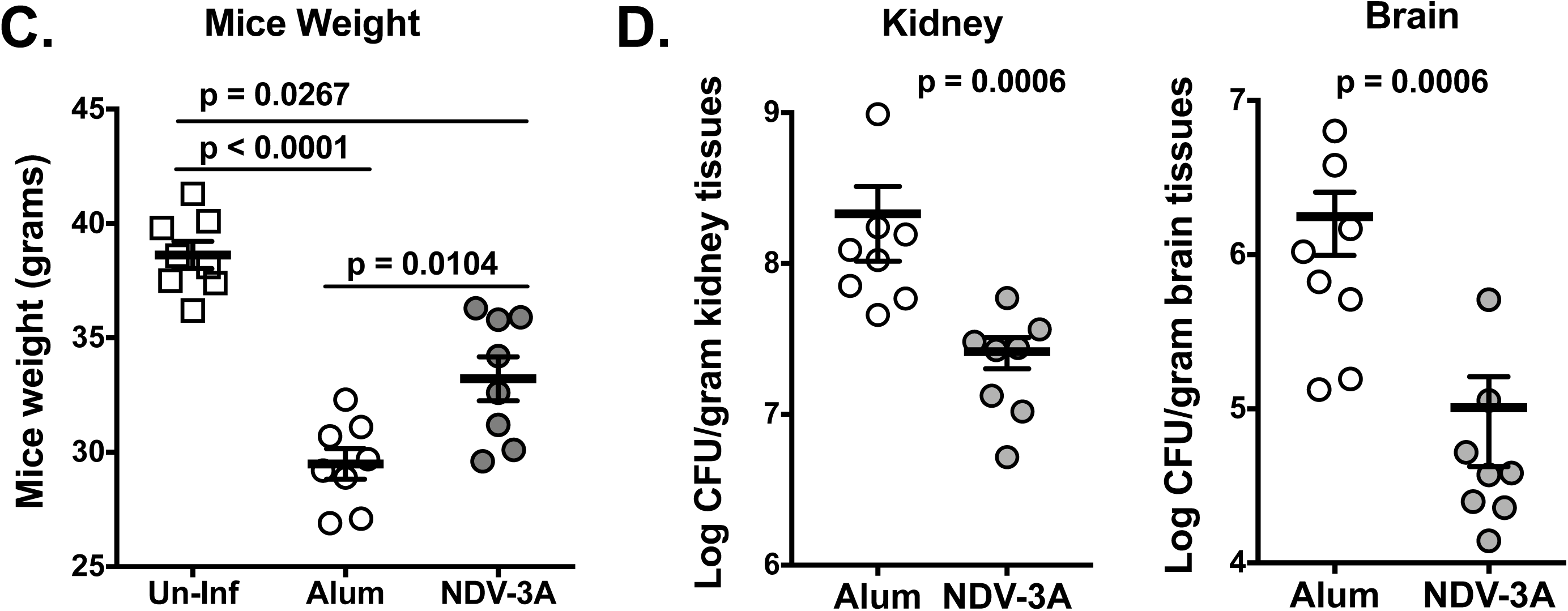
NDV-3A-vaccinated mice are protected from *C. auris* infection. **A)** Outbred CD-1 mice were immunized three times with either NDV-3A or alum alone. Neutropenia was induced by two doses of cyclophosphamide and cortisone acetate on day −2 and +3 of infection. After two weeks of final boost, the mice were infected through tail vein with lethal 5x10^7^ CFU/mouse dose. The mice from both groups were divided into two arms. One arm was observed for survival for 21 days, and another arm were sacrificed on day 4 post infection. **B)** NDV-3A vaccinated mice were protected from *C. auris* disseminated infection. The mice survival was compared by using Mantel-cox Log-rank Test. **C)** After 4 days post infection, NDV-3A or alum control vaccinated mice (n=8/group) were weighed sacrificed, and kidneys and brains were harvested and processed for fungal burden determination. Fungal burden was represented as CFU/g of organ weight. Statistical significance was determined by Mann-Whitney Test. P values <0.05 was considered significant.

Histopathological examination of tissues collected from mice sacrificed on day 4 post infection, confirmed inhibition of disseminated fungal infection in kidneys of NDV-3A-vaccinated mice, while alum-treated mice had multiple abscesses containing *C. auris* cells throughout the organ (Figure 6A). Compared to kidneys, the brain had an overall 100-fold reduced fungal burden (Figure 5D), which was also apparent in the histopathological images (Figure 6). While overall only a few cells could be visualized in the brain, alum-vaccinated mice appeared to have more fungi compared to brains of NDV-3A-vaccinated mice. Interestingly, in both cases, *C. auris* cells were localized in the blood capillaries, rather than the brain tissue itself (Figure 6B).

**Figure 6.**
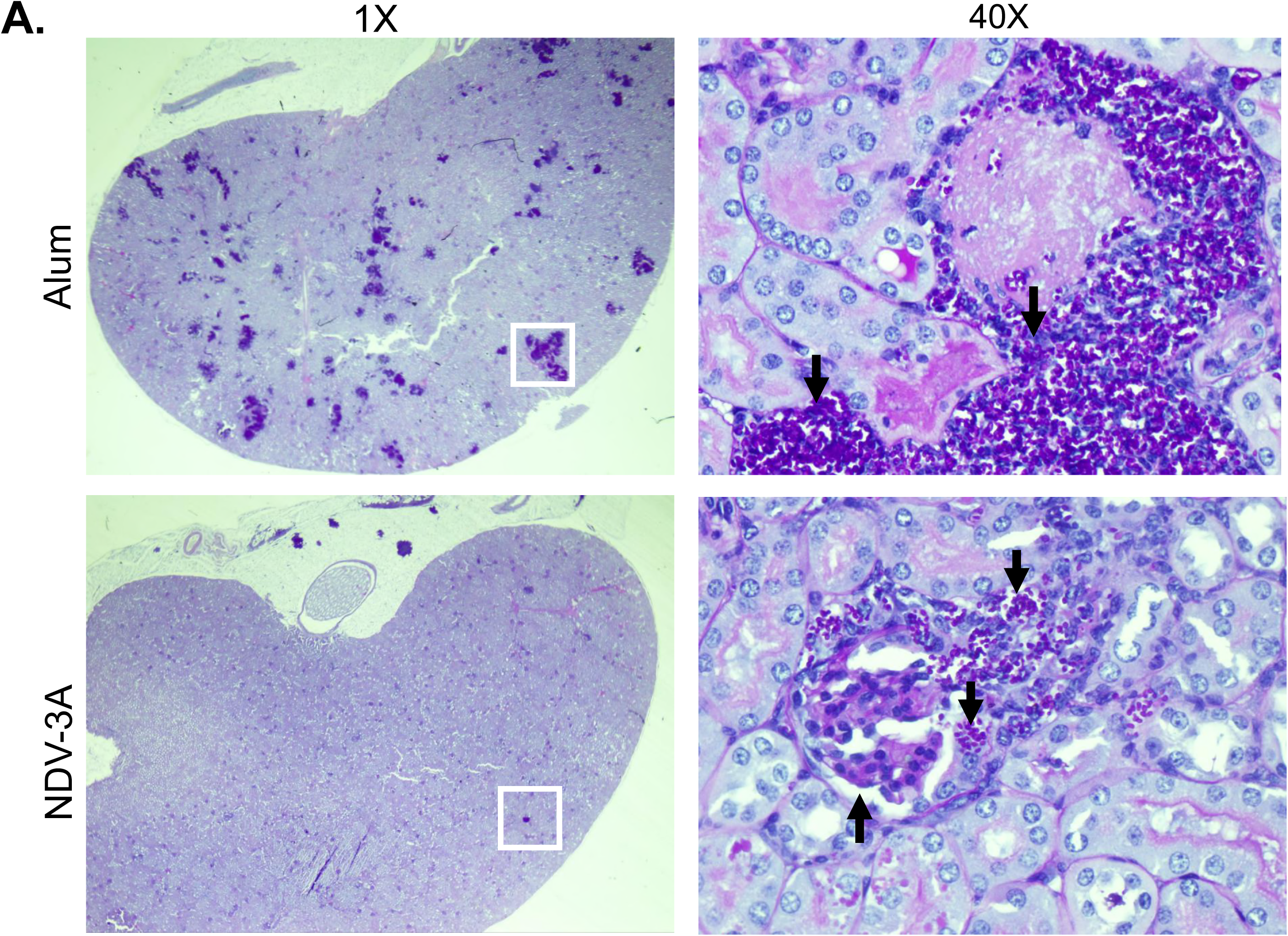
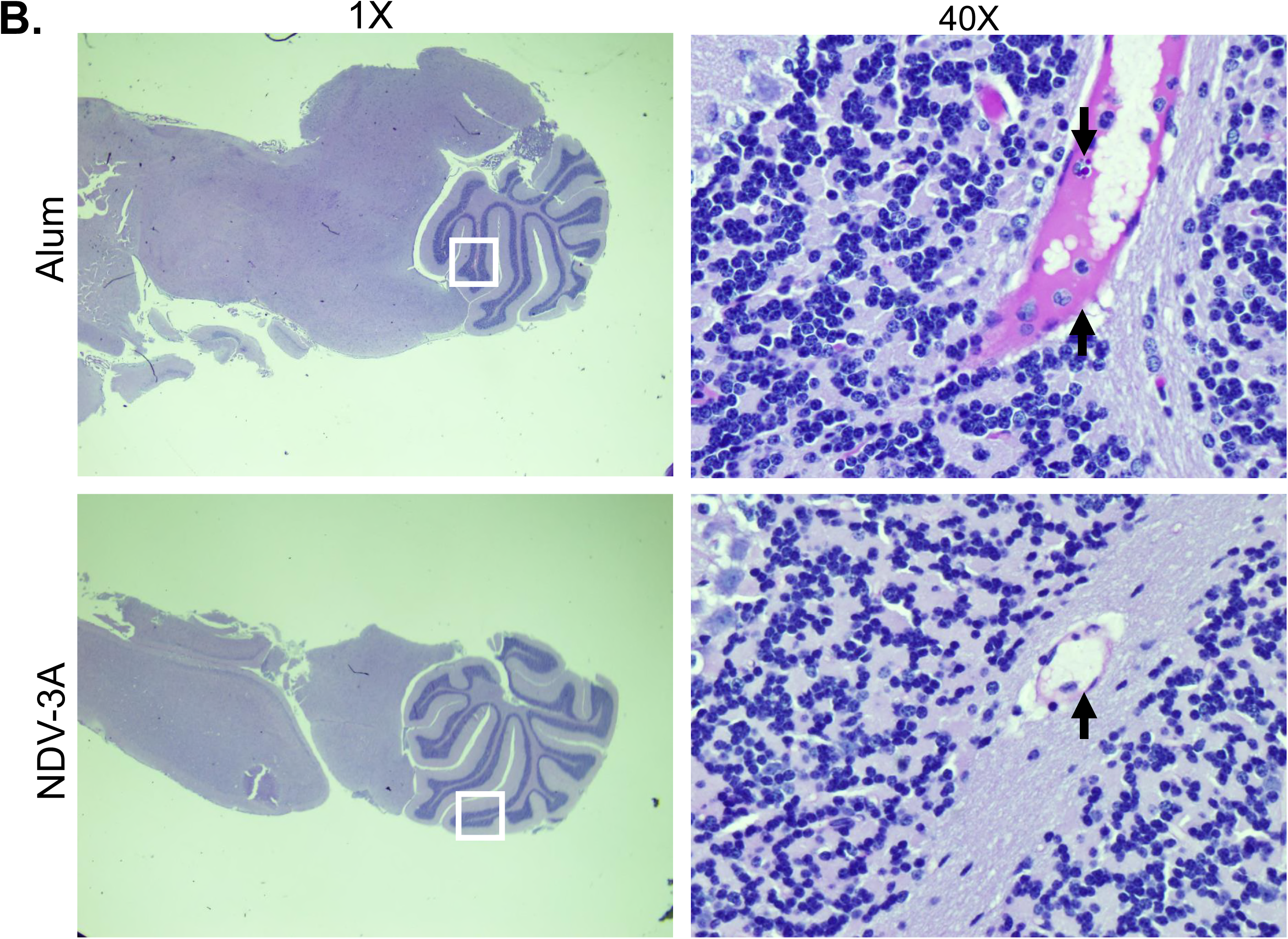
Comparison of histopathology of kidneys and brain tissue sections from NDV3-A- or alum-vaccinated and *C. auris* infected mice. Tissue sections obtained on 4 days after the infection were stained with PAS. **A) Kidneys.** The kidney from the control alum mouse had numerous abscesses with visible *C. auris* fungal cells throughout the tissue. **B) Brain.** NDV-3A-vaccinated mouse did not show any visible abscesses or *C. auris* organism but alum mouse brain was visibly infected with *C. auris*. The arrow facing down show the *C. auris* in the tissues, and arrows facing up show the blood capillaries.

### NDV-3A vaccination potentiated the activity of micafungin, against *C. auris* disseminated infection

In the clinical setting, any vaccine approach is likely to be used in combination with standard antifungal therapy. Given that NDV-3A vaccination provided significant protection against *C. auris* candidemia, we questioned if vaccination could potentiate the activity of antifungal drugs, *in vivo*. To investigate this possibility, we vaccinated mice with NDV-3A or alum as above, made them neutropenic and then infected them with *C. auris*. Twenty-four hours post infection, one group of NDV-3A-vaccinated or alum control mice were treated with sub-inhibitory concentrations of micafungin (0.5 mg/kg body since CAU-9 is micafungin sensitive [MIC=0.125 μg/ml]) daily for 1 week. The micafungin treatment regimen was determined in pilot studies to be moderately protective (data not shown). As expected, NDV-3A protected 40% of mice from succumbing to infection, while micafungin provided only 20% protection. Interestingly, when used in combination, NDV-3A and micafungin significantly enhanced the overall survival of mice to >70% (p=0.04, compared to NDV-3A-vaccination alone; and p=0.001, compared to micafungin alone) (Figure 7). These results show that NDV-3A vaccine acts additively with active antifungals in protecting mice from *C. auris* candidemia.

**Figure 7.**
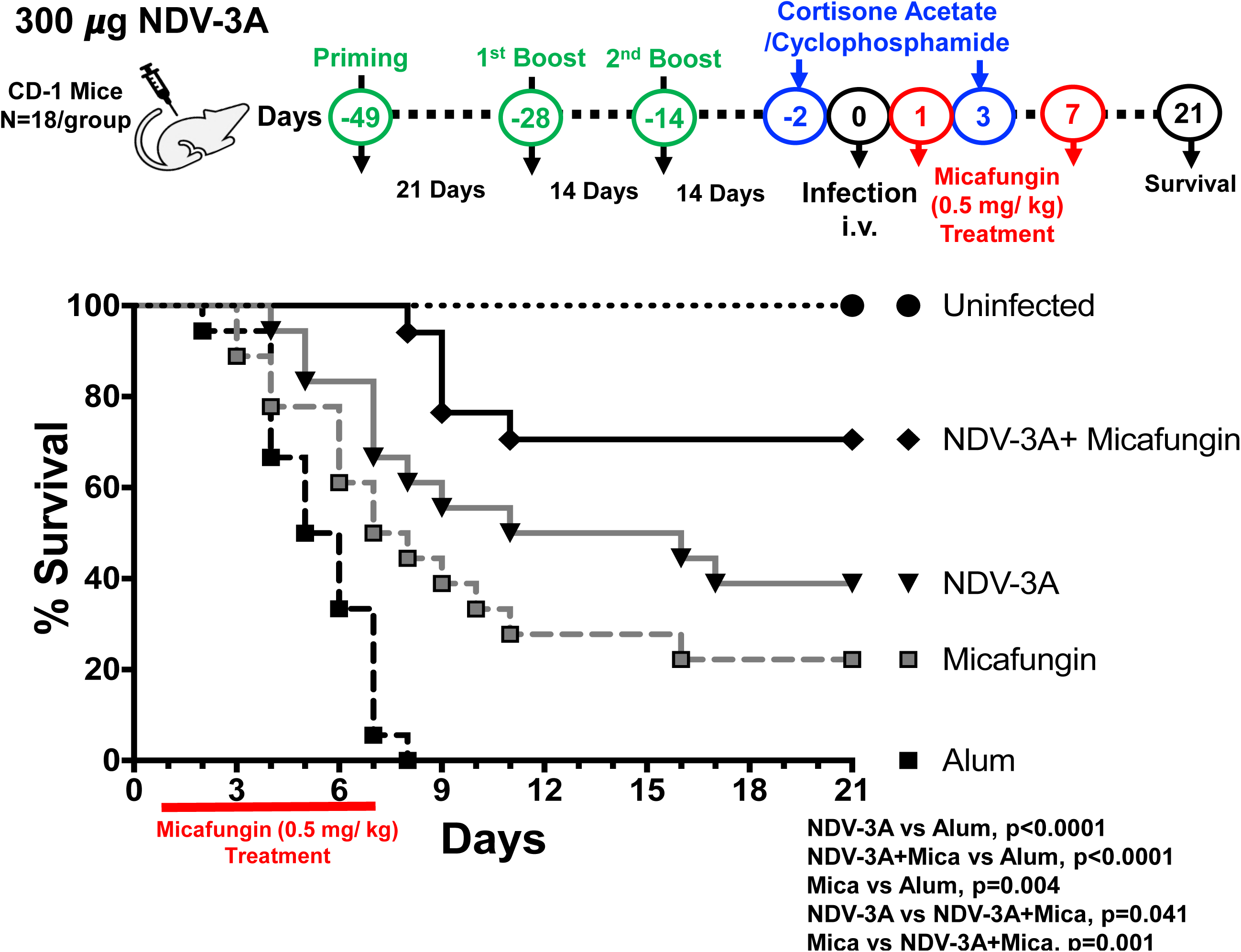
Combination of micafungin and NDV-3A vaccine additively protect neutropenic mice from *C. auris* bloodstream infection. NDV-3A- or alum control-vaccinated mice were made neutropenic and infected with *C. auris*. The mice were divided into four groups: NDV-3A vaccine alone, micafungin treatment alone, NDV-3A vaccine + micafungin, alum alone (n=18 from two independent experiments). The mice were evaluated for their survival for 21 days. Both NDV-3A vaccine and micafungin treatment group showed significantly higher survival compared to alum control group. The NDV-3A +micafungin treatment showed further enhanced survival compared to NDV-3A or micafungin treatment alone mice. The mice survival was compared by using Mantel-cox Log-rank Test. P values <0.05 was considered significant.

## Discussion

The simultaneous global emergence of the pathogenic yeast *C. auris*, its ability to persist in healthcare-settings, combined with its potential to resist all classes of antifungal drugs, has resulted in a renewed focus of the Centers for Disease Control (CDC), health-care professionals and scientists alike, on multidrug resistant fungal organisms [3, 8]. To add to the problem, *C. auris* reportedly possesses the capacity to adhere to and develop inherently drug-resistant biofilms on abiotic surfaces [10, 27], thus bolstering its ability to persevere in hospital environments, and cause outbreaks. Disseminated infections due to *C. auris* in susceptible patients (e.g. advanced age, presence of central venous catheter, surgery, prolonged hospitalization), are associated with unacceptably high mortality rates between 30-60% [6, 7]. It is unlikely that successful treatment will be achieved with antifungal drugs alone, even with proper antibiotic stewardship. Identification of alternative strategies to prevent and treat infections caused by this fungus are needed. Improved therapy is achievable by either discovering novel molecular targets in its genome, or identifying those that may be homologous to other therapeutically-targeted proteins from phylogenetically related fungi.

We applied computational molecular modeling and bioinformatic strategies to demonstrate that *C. auris* possesses homologs of *C. albicans* major adhesin/invasin protein, Als3p. This was an intriguing finding, since Als3p is a hyphae specific protein and *C. auris* does not form filaments. The presence of Als3p-homologs in *C. auris* adds to the list of other genes or gene products with homologies to those associated with virulence and drug resistance in *C. albicans*. For example, homologs of several *C. albicans* efflux genes belonging to the major facilitator superfamily (MFS), *MDR* and the ATP-binding cassette (ABC) transporter families have been identified, suggesting that efflux is likely a potential resistance mechanism mediating multidrug resistance in *C. auris* [28, 29]. In fact, homologous genes of *C. albicans* virulence proteins such as Serine/Threonine Enzyme-related proteins, mannosyl transferases, a number of secreted aspartyl proteases, as well as kinases involved in virulence and antifungal stress response such as Hog1 protein kinase, 2-component histidine kinase etc., have been identified in the *C. auris* genome [30]. Our discovery of Als3p-like proteins in *C. auris* clearly indicates that despite its extremely high genomic divergence from *C. albicans*, core gene families involved in adhesion/invasion, acquiring drug resistance and other virulence-related genes are conserved in *C. auris*; perhaps also a reason for its persistent nature and its success as a MDR organism.

Identifying Als3p homologs in *C. auris* makes it a target for vaccine strategies involving the NDV-3A vaccine, which was developed based on the N-terminus region of *C. albicans* Als3, and formulated with alum [18]. Our group has previously reported on the efficacy of NDV-3 (a His-tagged N-terminus of recombinant Als3p formulated with alum) in preventing disseminated candidiasis in mice infected with *C. albicans* [17]. Vaccination with NDV-3 also protects mice from hematogenously disseminated candidiasis due to other *Candida* species including *C. glabarata, C. tropicalis*, and *C. parapsilosis* (data not shown). NDV-3A (N-terminus of recombinant Als3p without his-tag and formulated with alum) was also shown to be safe when given to healthy volunteers [18]. In a Phase 1a/2b clinical trial, NDV-3A was deemed safe and immunogenic in women who suffer from recurrent vulvovaginal candidiasis and protected them from infection [19, 20]. Most recently we have described that anti-Als3p antibodies generated from NDV-3A-vaccinated patients (and not alum-treated) had the potential to affect properties of adherence, filamentation and biofilm formation in *C. albicans* [20]. Thus, we questioned if anti-Als3p antibodies could recognize *C. auris*, interfere in functions characteristically associated with Als3p, and prove clinically valuable as vaccine-based strategies against *C. auris*. Using different and complementary approaches (immunofluorescence, flow cytometry, and cell-based ELISA), we validated that NDV-3A vaccination-generated high titer anti-Als3p antibodies that recognized different strains of *C. auris* (Figure 2, 3A). Importantly, the endpoint IgG antibody titer determined by regular Als3p antigen ELISA and *C. albicans* cell-based ELISA was the same. This finding clearly demonstrates that the two assays are comparable and cell-based ELISA can correctly estimate the antigen-specific antibody titer in serum. Based on these results, we can conclude that although *C. auris* cross-reactive IgG antibodies were 4-fold lower than the Als3p antigen specific IgG antibodies, these titers were still sufficiently high (Figure 3A).

The protective efficacy of NDV-3A vaccine against *C. albicans* infection has been correlated with high antibody titers and Th1/Th17 cell mediated immune responses [17, 21, 23]. An effective immunity against a pathogen is usually comprised of both antibodies and memory T cell immune responses. Therefore, we examined the splenocytes of NDV-3A-vaccinated mice and detected high levels of *C. auris* cross-reactive CD4+ T cells responses in NDV-3A-vaccinated mice, which was similar to Als3p-specific T cells in magnitude. Further, these CD4+ T cells responses (both *C. auris* and Als3p-specific) were marginally shifted towards a Th2-type, probably because alum is known to favor Th2 cell induction [31, 32]. Further, the NDV-3A vaccination also induced high frequency of Als3p-specific Th17 cells that cross-reacted to heat-killed *C. auris*. These results are in line with our previous reports showing that NDV3 and NDV-3A vaccines induce strong Als3p-specific Th17 immune responses [18, 19, 21-23]. Together, these results highlight that NDV-3A vaccine induced robust *C. auris* cross-reactive antibody and T cell immune responses (Figure 3).

The contribution of anti-Als3p antibodies towards abrogation of virulence potential of *C. auris* was illustrated effectively by its ability to block *C. auris* biofilm formation, *in vitro* (Figure 4A). Additionally, opsonization of *C. auris* with anti-Als3p antibody containing sera from NDV-3A-vaccinated mice resulted in efficient immune recognition of the pathogen by mice macrophages, leading to increased opsonophagocytic killing (Figure 4B), a feature that was noticed with the activity of anti-Als3p antibodies and *C. albicans* [20, 33]. This begged the potential of evaluating NDV-3A for protection against *C. auris* infections, *in vivo*. Indeed, the results from *in vitro* assays translated successfully *in vivo*, where prophylactic vaccination by NDV-3A protected some neutropenic mice from disseminated candidiasis caused by *C. auris* (Figure 5B). Thus, NDV-3A vaccine can be used as a potential additional strategy targeting *C. auris* infections even in the setting of immunosuppression. The exact mechanism behind this protection is yet to be identified but it is possible that the produced anti-Als3p antibodies neutralized the function of the Als3p-like protein which might be required for *C. auris* virulence. Furthermore, although the use of cyclophosphamide in mice results in leukopenia, there is evidence that its effect on tissue macrophages is less severe. In fact, deficiency in alveolar macrophages due to continuous administration of cyclophosphamide was more of a chronic and gradual reduction that never reaches complete deficiency [34]. Thus, the residual tissue macrophages in the cyclophosphamide/cortisone acetate could contribute to the protection seen *in vivo* given the fact that anti-Als3p antibodies enhance OPK activity of macrophages *in vitro* (Figure 4B). Of great clinical relevance is the additive effect in protecting mice from *C. auris* infection when the vaccine was given with micafungin (Figure 7). This result provides a strong rationale for the combined use of active NDV-3A vaccination with antifungal drugs in multidrug-resistant lethal *C. auris* infections and gives hope to improved treatment outcomes.

In summary, the unique property of *C. auris* to persist outside human body in the harsh environmental settings of the hospital, its rapid evolution of drug resistance, and its propensity to infect immunosuppressed hospitalized individuals, increasingly threaten global and personal health. The identification of Als3p homologs in *C. auris* opens an avenue for novel immunotherapeutic approaches utilizing either the currently available NDV-3A vaccine, or perhaps an anti-Als3p antibody mediated passive vaccination strategy in the near future. These immunotherapeutic approaches could enhance successful treatment of infections caused by such fungal “superbugs”, thereby reducing morbidity and mortality. Given the safe profile of NDV-3A in humans, future studies will focus on testing this vaccine in patients at high risk of acquiring *C. auris* infection (e.g. colonized patients) and/or those who already suffer from the infection (e.g. therapeutic vaccine combined with antifungal agents).

## Materials and Methods

### Organisms and culture conditions

The *C. auris* strains (CAU-01, East Asian clade, ear; CAU-03, African clade, blood; CAU-05, South American clade, blood; CAU-07, South Asian clade, blood; and CAU-09, South Asian clade, bronchoalveolar lavage [BAL]) were obtained from Dr. Shawn Lockhart at Centers for Disease Control and Prevention (CDC, Atlanta). For routine culturing, *C. albicans* (SC5314) and *C. auris* were grown in Yeast Extract Peptone Dextrose (YPD) broth overnight at 30°C with shaking at 200 rpm. Cells were washed with 1x phosphate buffered saline without Ca^++^/Mg^++^ (PBS, Gibco by Life Technologies) three times prior to counting blastopores with a hemocytometer. For germinating *C. albicans*, 5x10^6^ yeast cells/ml in RPMI1640 media (supplemented with L-Glutamine) were allowed to form germ tubes at 37°C for 75 minutes with shaking at 200 rpm. For staining, cell based ELISA, and splenocytes stimulation, the *C. auris* was grown in above germination conditions. For biofilm formation assay, the cells were incubated at 37°C in yeast nitrogen base (YNB) medium for 24 hours.

For heat killing of yeast cells, 5 x 10^6^ cells /mL of PBS were incubated at 65°C for 45 minutes. Yeast cell death was confirmed by plating the heat-subjected cells on fresh plate and incubating the cells at 30°C for several days. For fixation, the yeast cells were incubated with 4% paraformaldehyde solution (in PBS) at 4°C for 1 hour.

### Sequence alignment and analysis

The *C. albicans* N-terminal <u>A</u>gglutinin-<u>L</u>ike <u>S</u>equence <u>p</u>rotein (Als3p, Gene Bank ID: AOW31402.1) amino acid sequence was aligned with the *C. auris* using protein BLAST and CLUSTAL-W (NCBI). The predicted Als3p homologs in *C. auris* proteome were further screened for homology among each other using CLUSTAL-W and Als3p homologs having >95% homology with each other considered as same protein. The number of Als3p amino acids showing similarities with *C. auris* proteins was represented as percent positive. The functional domains were identified by using different online bioinformatics tools for GPI-anchor (http://gpi.unibe.ch/), amyloid sequence (http://protein.bio.unipd.it/pasta2/), and Ser/Thr rich sequence (https://www.ebi.ac.uk/Tools/pfa/radar/) prediction. The 3-D protein structure models were built using amino acid sequences and the templates available in the Swiss-model database [35-37] (https://swissmodel.expasy.org/). Briefly, the templates were searched in SWISS-MODEL template library (SMTL) using BLAST and HHBlits. The target sequence was searched with BLAST against the primary amino acid sequence contained in the SMTL. The target-template alignment was performed to build the model by using ProMod3, and coordinates that were conserved between the target and the template were copied from the template to the model. Insertions and deletions were re-modelled using a fragment library, and the side chains were rebuilt. Finally, the geometry of the resulting model is regularized by using a force field. In case loop modelling with ProMod3 fails, an alternative model is built with PROMOD-II. The models showing high accuracy values were finalized for similarity comparisons.

### Immunization of mice

We used 4-6 week old outbred ICR (CD-1) mice in this study. The NDV-3A vaccine was formulated by mixing 300 μg of *C. albicans* recombinant Als3p with 200 μg alum adjuvant per dose. The recombinant Als3p was produced in *Saccharomyces cerevisiae*, and was a Gift from NovaDigm therapeutics. For immunization, the NDV-3A vaccine or alum was injected into mice subcutaneously (s.c.) at day 0, 21 and 35. Mice were bled 14 days after final immunization and serum was isolated for antibody titer determination.

### Confocal microscopy and flow cytometry

The yeast or germ tube cells (2x10^6^ cell) from either *C. auris* or *C. albicans* were fixed with 4% paraformaldehyde at 4°C for 30 minutes. After blocking the cells with 3% bovine albumin serum solution (in 1x PBS), these cells were added to 96-well plate and centrifuged to pellet the cells. The pellet was resuspended in 100 μl of anti-NDV-3A or alum serum diluted at 1:500 in 1x PBS and incubated for 1 hour at room temperature. The cells were washed three times with 1x PBS prior to adding 100 μl of Alexa Fluor 488 labelled anti-mouse IgG detection antibodies (1:100 dilution in PBS). After 1 hour of incubation at room temperature, the cells were resuspended in 300 μl of PBS and analyzed using confocal microscopy or flow cytometry.

For confocal microscopy, 20 μl of stained cells were added to the glass slides and covered with cover slips. The images were taken at 40x resolution using Laser Scanning Confocal microscopy (Leica Confocal Microsystem). For flow cytometry, the stained cell suspension was transferred to flow tubes and 20,000 events were acquired using LSR II flow cytometer (BD Biosciences). Flow cytometry data were analyzed using FlowJo software (Version 10).

### Elisa

Ninty six-well plates were coated with 10 μg/ml of Als3p in bicarbonate/carbonate coating buffer (pH 9.6) overnight at 4°C. The next day, the plates were washed three times with 1x wash buffer (PBS containing 0.05% tween-20) and blocked with 3% BSA solution for 2 hours at room temperature. After washing three times, diluted serum samples were added to the plates in duplicates and incubated for two hours. After incubation, the plates were washed three times and 1:1000 diluted anti-mouse IgG antibodies (Jackson, Cat#115-035-164) labelled with peroxidase were added and incubated for 1 hour at room temperature. Finally, the plates were washed five times with washing buffer, TMB (3,3′,5,5′-Tetramethylbenzidine) substrate (Invitrogen, Cat#00-4201-56) was added. Color development was allowed for 5-10 minutes and absorbance was measured at 450 nm after the reaction was stopped with 1 N sulfuric acid (Sigma, Cat#339741).

To determine the *C. auris* cross-reactive antibodies in NDV-3A vaccinated mice, we used a cell-based ELISA. As above, germinated *C. albicans* (SC5314) or *C. auris* (CAU-09) were counted and fixed in 4% cold paraformaldehyde (Sigma-Aldrich, Cat#158127) for 30 min. The fixed cells were washed with 1x PBS. The U-bottom 96-well plates were blocked overnight with 1x blocking solution (3% BSA in 1x PBS, Thermo Fischer), and next day 2-fold serial dilutions of 100 μl serum samples (diluted in 1x blocking solution) per well were added in duplicates. Further, a total of 10^7^ fixed cells/well/100 μl of *C. auris* and germinated *C. albicans* were added to these serum sample-containing wells. The plates were incubated for 2 hours at room temperature. After incubation, the cells in the 96-well plate were washed three times with 1x PBS and secondary HRP-conjugated anti-mouse IgG antibodies were added as per the manufacturer instructions. After 1 hour incubation at room temperature, the cells in 96-well plate were washed with 1x PBS. Further, 100 μl/well TMB substrate was added to each well and incubated until the color developed (normally 5-20 min). The color reaction was stopped by adding 50 μl 1N H_2_SO_4_ per well and plates were centrifuged at maximum speed to pellet the cells. One hundred microliter colored substrate supernatant from each well were transferred to fresh flat bottom 96-well plates and OD was measured at 450 nm. The endpoint titers in NDV-3A vaccinated mice were determined by plotting mean OD450 vs serial serum dilution, and noting the highest dilution with significantly higher OD450 compared to alum vaccinated mice group.

### Intracellular cytokine staining and flow cytometry

The splenocytes were harvested from the NDV-3A or alum immunized mice by homogenizing individual spleens in 100 μm cell strainer. The RBCs were lysed by 1x RBC lysis buffer (Santa Cruz Biotech, Dallas, Cat# SC-296258), and filtered through the 100 μm sterile filters. The cells were resuspended in 1x RPMI supplemented with 10% FBS, counted and plated at 1x10^6^ splenocytes/ 100 μl/ well in a U-bottom 96-well plate. The splenocytes were kept unstimulated or stimulated with 100 μl/well of 10 μg/ml recombinant Als3p, or 3x10^6^ cells/well heat-killed *C. auris* for 5 days at 37°C. On day 5, 100 gl of the culture supernatant was replaced with fresh RPMI media containing 50 ng/ml PMA (phorbol 12-myristate 13-acetate, Sigma, Cat# P8139), and 500 ng/ml ionomycin (Sigma, Cat#I3909). After 1 hour, protein transport inhibitor cocktail (eBioscience, Cat# 00-4980-03) was added to each well at final 1x concentration, and plates were incubated for another 3 hours. Next, the cells were stained with extracellular CD3 APC-Cy7 (BD Biosciences, Cat#557596) and CD4 PerCP Cy5.5 (BD Biosciences, Cat#550954) antibodies (0.25 μg/sample) for 30 minutes at room temperature. Subsequently, the cells were washed, fixed and permeabilized with Cytofix/Cytoperm solution (Invitrogen, Cat#00-5123-43) and then stained intracellularly with IFN-γ APC (BD Pharmingen, Cat#554413), IL-4 FITC (eBioscience, Cat#11-7042-82) and IL-17 PE (R&D Systems, Cat#IC7211P) antibodies (0.25 μg/sample) for 30 minutes at room temperature. The stained cells were acquired in BD LSR-II flow cytometer (BD Biosciences), and at least 30,000 CD3+ T cells were recorded. The data was analyzed using FlowJo (Version 10) software.

### Biofilm formation assay

Biofilms were developed in 96-well polystyrene microtiter plates as previously described, with slight modifications [38]. Briefly, 95 μL of *C. auris* cells (2 × 10^5^ cells/ml in YNB medium) was added to the wells containing 5 μL of test or control mouse serum (5% serum vol/vol), and incubated at 37°C. Control wells had no serum. After 24 hours, wells were washed twice with PBS, and the extent of biofilm formation was quantified by XTT assay (490 nm) [38].

### Opsonophagocytic killing (OPK) assay

The opsonophagocytic killing assay was based on a modification of a previously used method [39]. *C. auris* yeast cells (CAU-09) were added into 96-well microtiter plates. Murine RAW 264.7 macrophage cells (American Type Culture Collection [ATCC# TIB-71], Rockville, MD) were cultured at 37°C in 5% CO_2_ in RPMI-1640 (Irvine Scientific, Santa Ana, CA) with 10% fetal bovine serum (FBS), 1% penicillin, streptomycin, and glutamine (Gemini BioProducts), and 50 mM β-mercaptoethanol (Sigma-Aldrich, St. Louis, MO). RAW 274.7 cells were activated by exposure to 1 ng/ml LPS (Sigma-Aldrich) for 24 hours. Activated RAW 264.7 macrophages were harvested after scraping with BD Falcon cell scrapers (Fischer Scientific) and added to the microtiter wells at a 1: 1 ratio of macrophages to *C. auris*. After 2 hours of incubation with gentle shaking, aliquots from the wells were quantitatively plated in YPD agar plates. The percent killing of *C. auris* was calculated using the following formula: 1-[CFUs from tubes with (mouse serum + *C. auris* + macrophages) /average CFU in tubes with (C. *auris* + macrophage)].

### Mice infection and treatment

CD-1 mice vaccinated with NDV-3A vaccine were infected with *C. auris* CAU-09 to evaluate the efficacy of the vaccine. Briefly, twelve days after the final boost, mice were made neutropenic by 200 mg/Kg cyclophosphamide delivered intraperitoneally (i.p.) and 250 mg/kg cortisone acetate (s.c.) administered on days −2 and +3, relative to infection. To prevent bacterial superinfection in the immunosuppressed mice, we added enrofloxacin (at 50 μg/ml) to the drinking water. Mice were infected through tail vein injection with 5x10^7^ CFU/mouse. For combination with antifungals, alum- or NDV3-A-vaccinated and infected mice were treated with a minimal protective dose of 0.5 mg/kg/day of the clinically used micafungin by i.p. administration. Treatment started after 24 hours of infection and continued until day +7. Mice were monitored for their survival for 21 days after the infection.

For fungal burden determination, mice were vaccinated, made neutropenic and then infected as above and then euthanized on day 4 post infection to collect kidneys and brain. The organs from each mouse were weighed, homogenized and quantitatively cultured by 10-fold serial dilutions on YPD plates. Plates were incubated on 37°C for 48 hours prior to enumerating colony forming units (CFUs)/gram of tissue. Finally, histopathological examination of kidneys or brain from mice sacrificed on Day 4 post infection, were fixed in 10% zinc-buffered formalin, embedded in paraffin, sectioned, and stained with Pacific Acid Schiff (PAS) stain.

### Ethical Statement

All procedures involving mice were approved by IACUC of Los Angeles Biomedical Research Institute (protocol 11672), according to the NIH guidelines for animal housing and care. Moribund mice according to detailed and well characterized criteria were euthanized by pentobarbital overdose, followed by cervical dislocation.

## Supplementary figures

**Figure S1:**
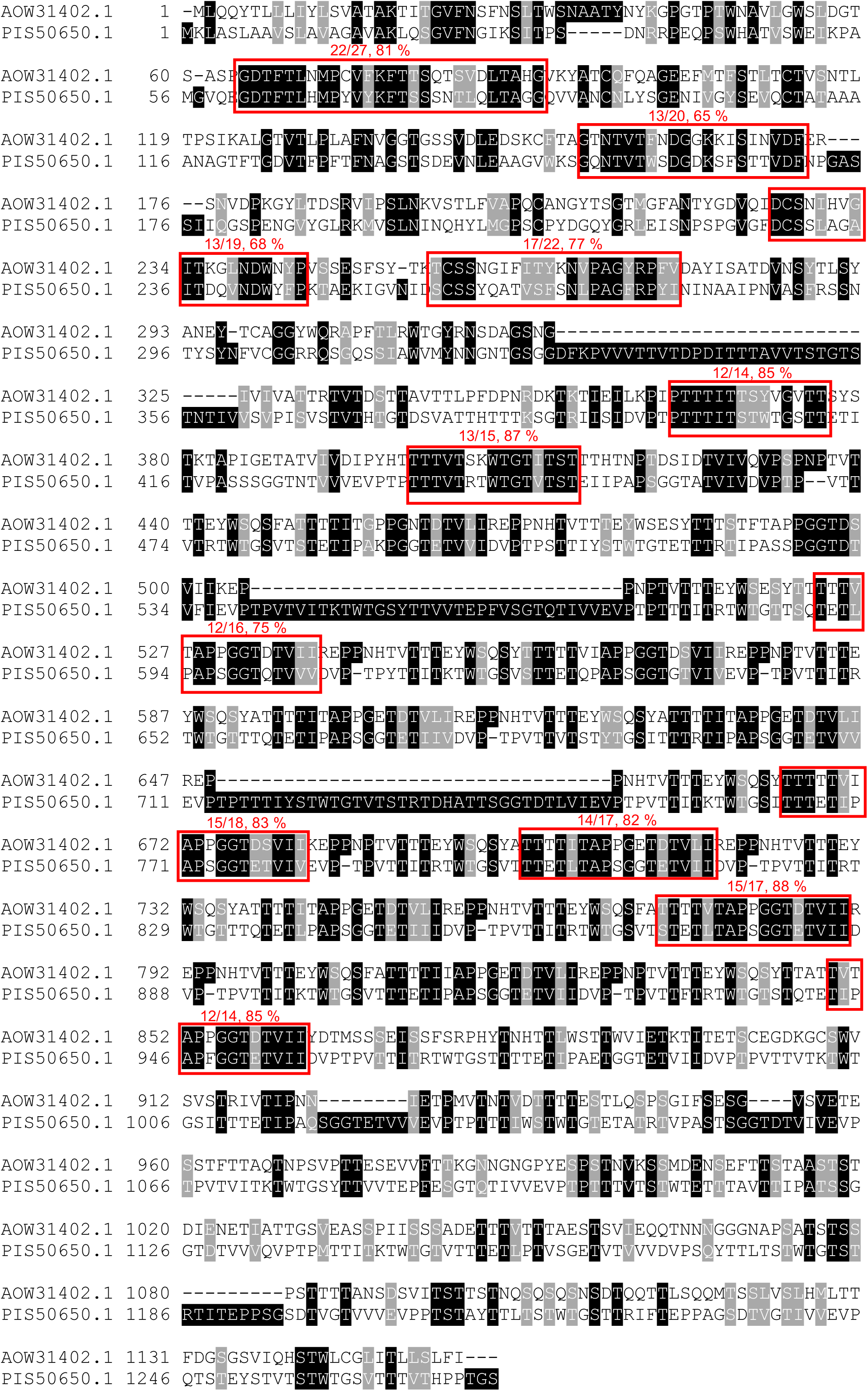
Protein sequence alignment between *C. albicans* Als3p (GenBank: AOW31402.1) and its homolog on *C. auris* (GenBank: PIS50650.1) using CLUSTAL-W.

**Figure S2:**
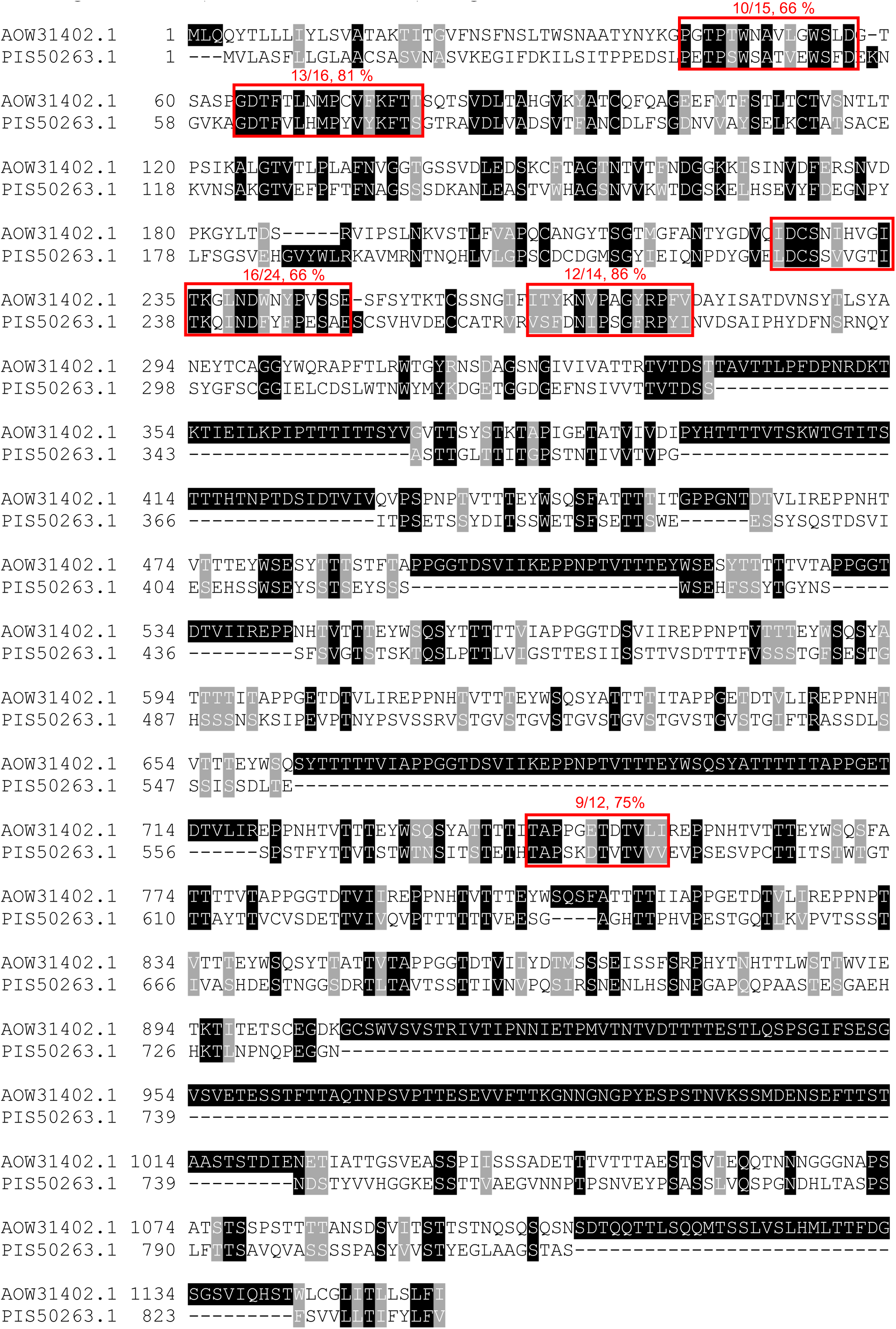
Protein sequence alignment between *C. albicans* Als3p (GenBank: AOW31402.1) and its homolog on *C. auris* (GenBank: PIS50263.1) using CLUSTAL-W.

**Figure S3:**
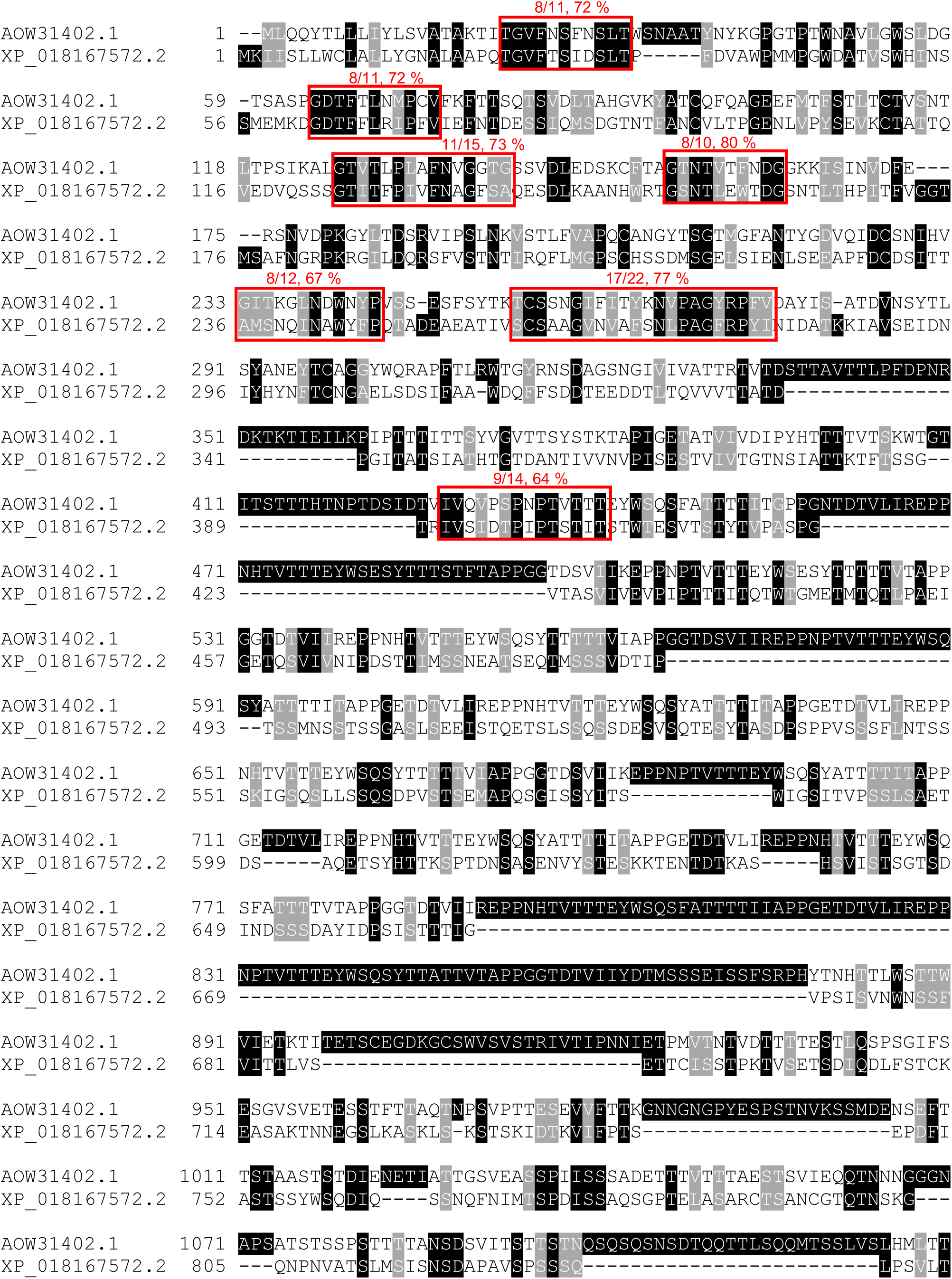
Protein sequence alignment between *C. albicans* Als3p (GenBank: AOW31402.1) and its homolog on *C. auris* (GenBank: XP_018167572.2) using CLUSTAL-W.

**Figure S4:**
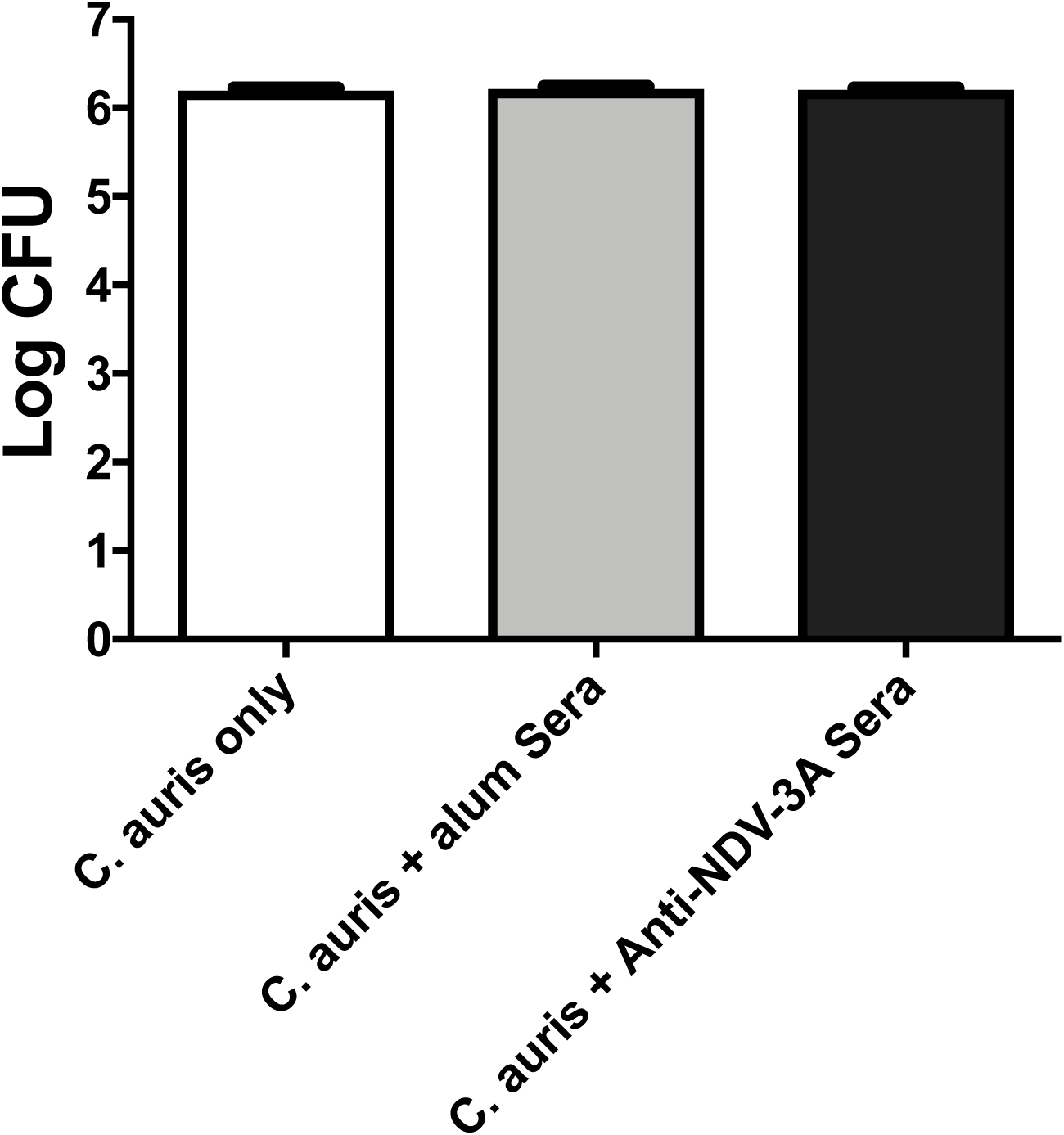
Sera from NDV-3A or Alum vaccinated mice have no effect on viability of *C. auris*

**Figure S5:**
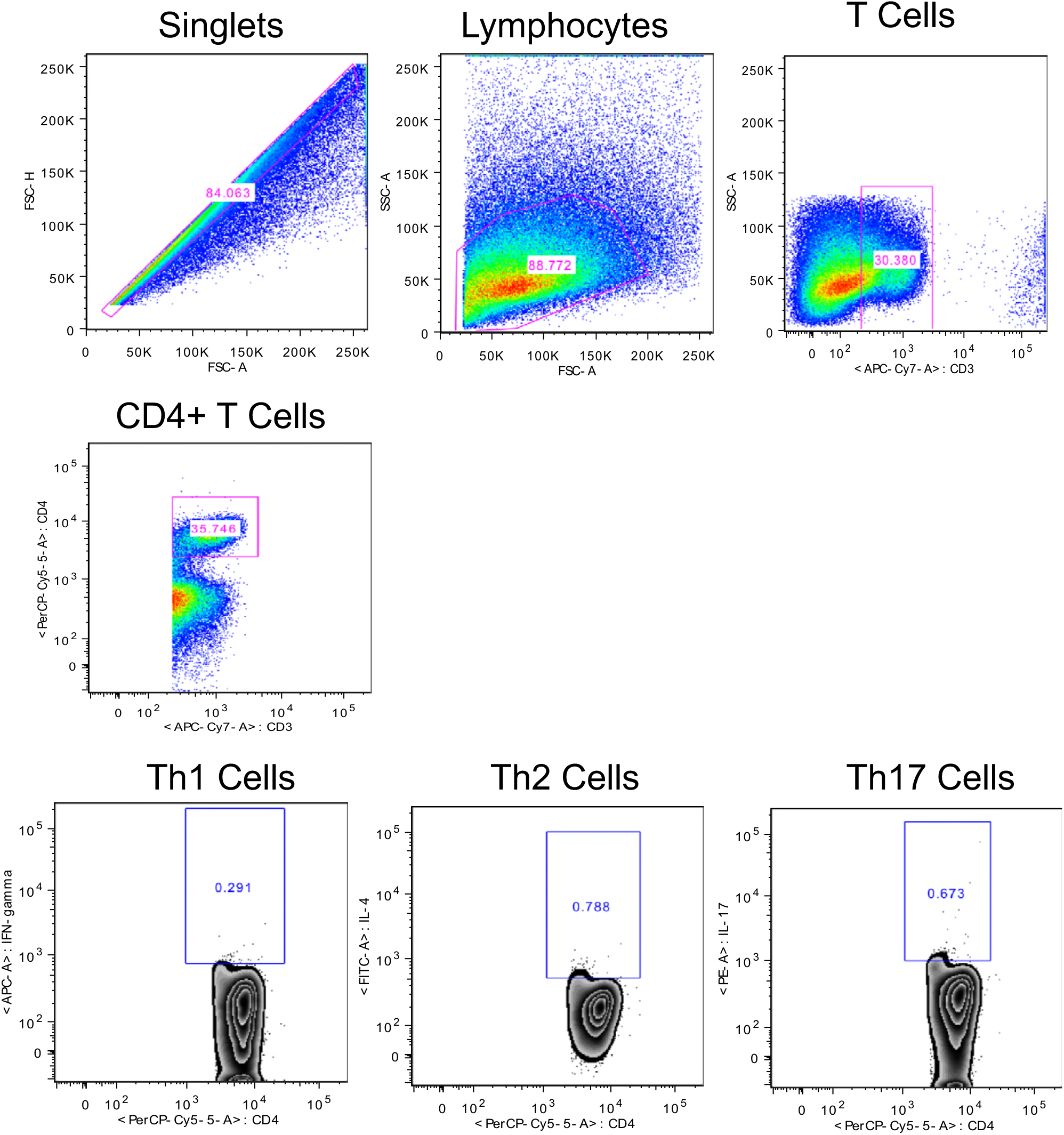
Intracellular cytokine staining gating strategy and Th1 (CD3+CD4+IFN-γ +), Th2 (CD3+CD4+IL4+) and Th17 (CD3+CD4+IL17+ phenotype determination.

**Figure S6:**
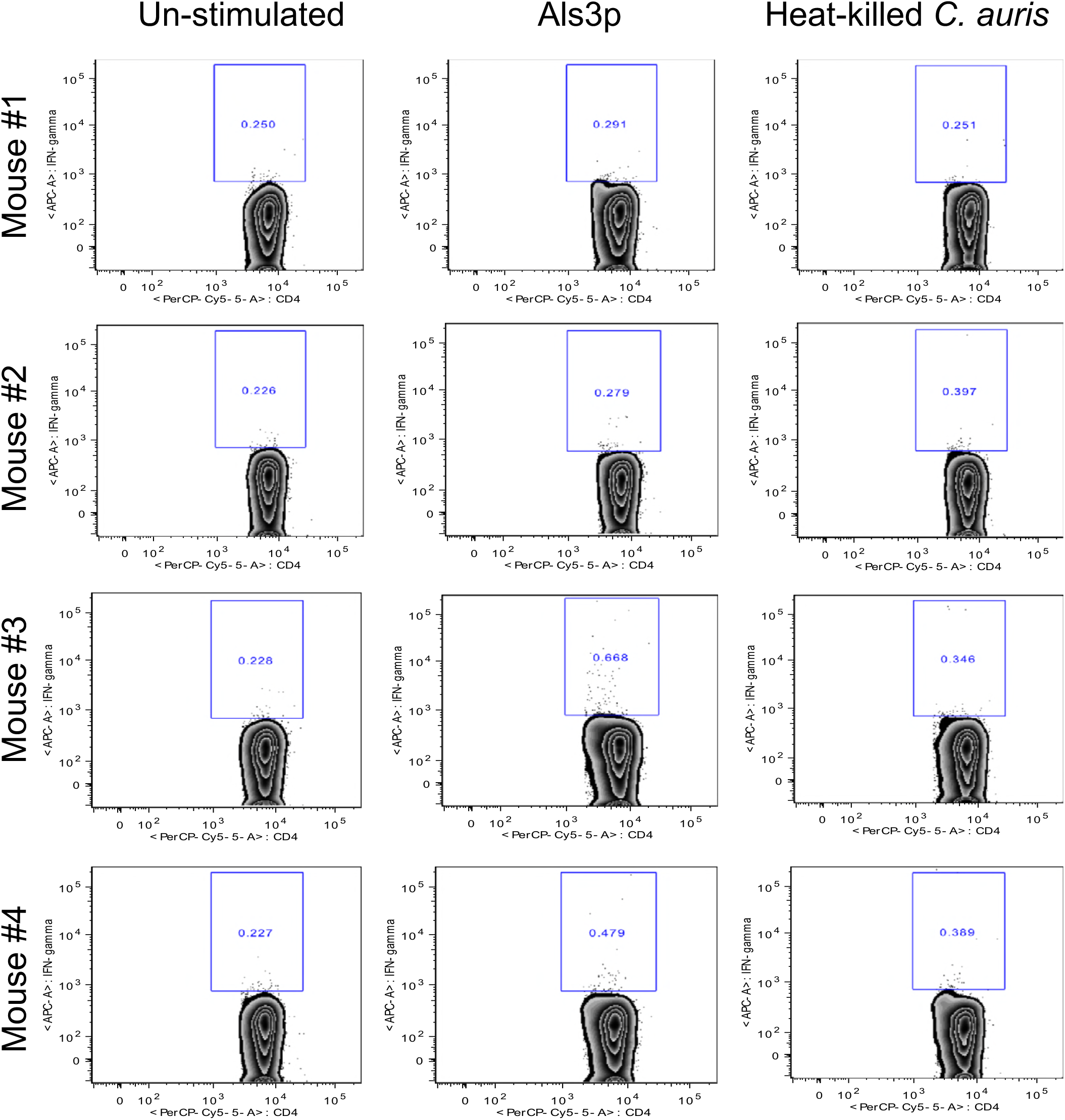
*C. auris* cross-reactive Th1 (CD4+ IFN-gamma+) cells in NDV-3A vaccinated mice

**Figure S7:**
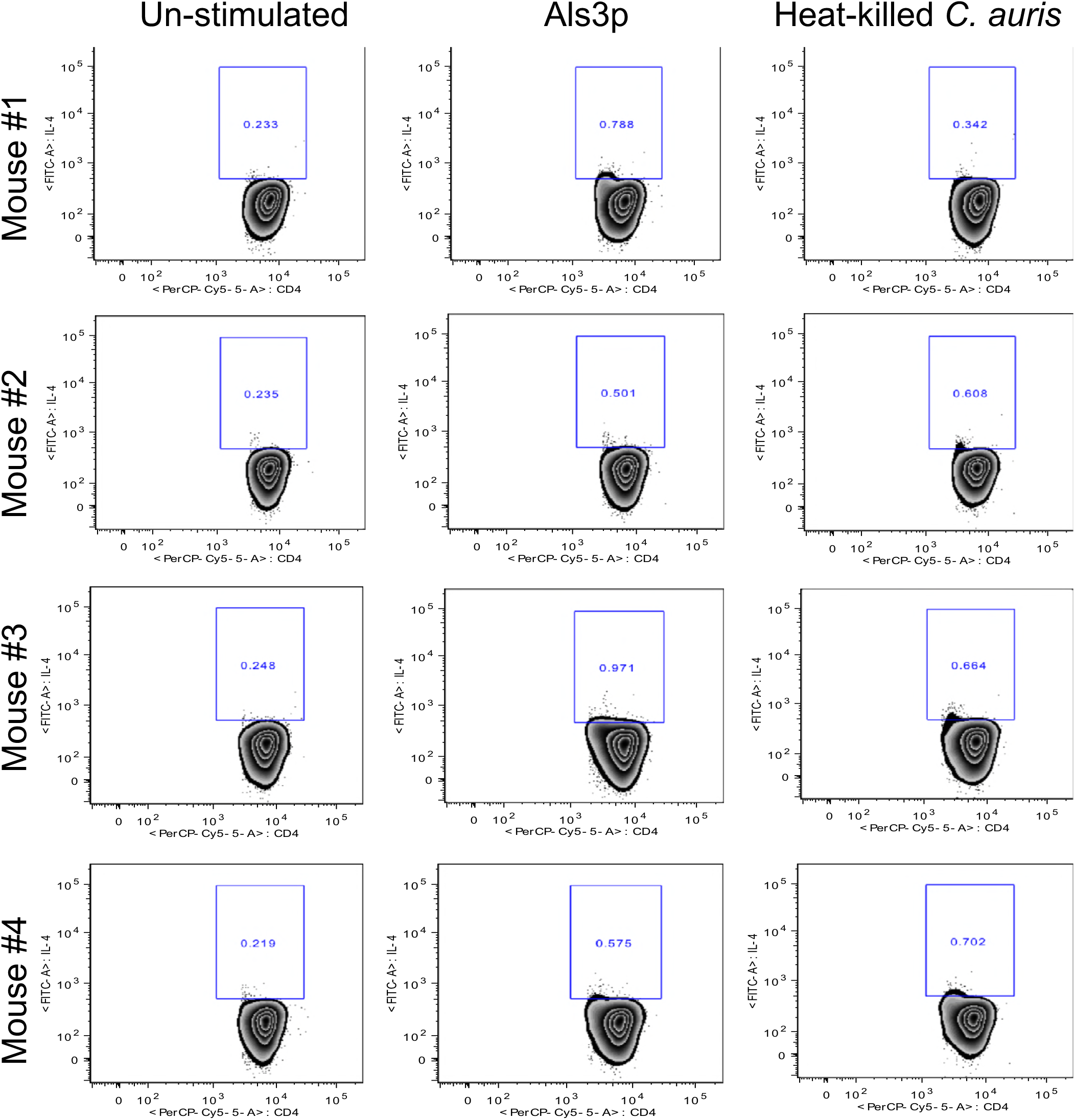
*C. auris* cross-reactive Th2 (CD4+ IL4+) cells in NDV-3A vaccinated mice

**Figure S8:**
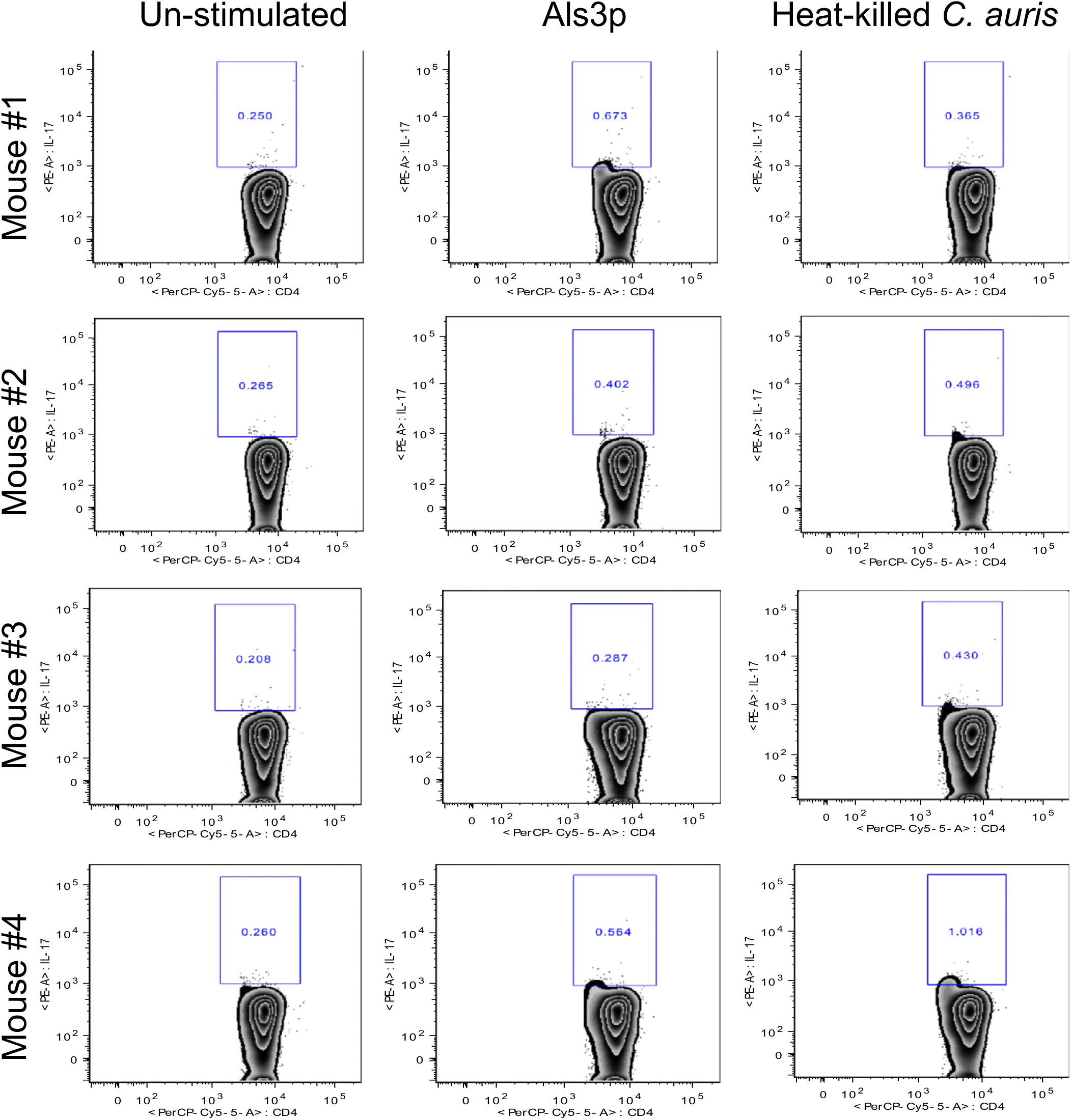
*C. auris* cross-reactive Th17 (CD4+ IL17+) cells in NDV-3A vaccinated mice

